# Common tissue-specific expressions and regulatory factors of c-KIT isoforms with and without GNNK and GNSK sequences across five mammals

**DOI:** 10.1101/2025.08.25.672062

**Authors:** Rikuto Goto, Naoaki Sakamoto, Akinori Awazu

**Affiliations:** Graduate School of Integrated Sciences for Life, Hiroshima University, Higashihiroshima, Hiroshima, Japan; Research Center for the Mathematics on Chromatin Live Dynamics, Hiroshima University, Higashihiroshima, Hiroshima, Japan

**Keywords:** c-KIT, RNA-binding proteins, isoforms, transcriptome data, central nervous system

## Abstract

c-KIT is a transmembrane receptor tyrosine kinase involved in various signaling pathways. Alternative pre-mRNA splicing of *KIT* results in isoforms that differ in the presence or absence of four amino acid sequences in the extracellular juxtamembrane region, such as, isoforms with and without the GNNK sequence (GNNK+ and GNNK−, respectively) in humans and mice, and those with and without GNSK (GNSK+ and GNSK−, respectively) in domestic dogs, cats, and sheep. These isoforms have been extensively studied as disease-associated (particularly tumors or cancer) splice variants with differing kinase activities. However, the expression patterns and regulatory factors of each isoform in various animal species without tumors or cancer remain poorly understood. Studying these aspects can provide the basis for understanding the associations between c-KIT isoforms and disease. Therefore, in the present study, a comprehensive expression analysis of c-KIT isoforms was conducted using tissue-wide transcriptome data from humans, mice, dogs, cats, and sheep. We found that the expression ratio of c-KIT isoforms differs across tissues, and such features are conserved across animal species: GNNK+ and GNSK+ isoforms have high expression ratios in the central nervous system, while GNNK− and GNSK− predominate in other tissues. Furthermore, *NOVA2*, *RBFOX1*, *RBFOX3*, and *DYRK1A* were suggested to be candidate factors regulating the selection of the alternative 5′ splice donor site of *KIT*.

## Introduction

c-KIT is the transmembrane receptor tyrosine kinase encoded by the *KIT* gene, which is activated by its ligand, stem cell factor (SCF), to trigger multiple downstream signaling cascades, such as PI3K, Src, JAK/STAT, PLC-γ, and MAPK pathways, essential for hematopoiesis, stem cell maintenance, gametogenesis, melanogenesis, mast cell development, and cell migration [1,2]. c-KIT comprises five N-terminal immunoglobulin-like domains, a single transmembrane domain, and C-terminal intracellular tyrosine kinase domains. In humans and mice, the extracellular juxtamembrane region, which is located adjacent to the fifth immunoglobulin-like domain (D5) that mediates dimer contacts [3], contains a four - amino-acid sequence Gly-Asn-Asn-Lys (GNNK) (Fig 1A and 1B). In domestic dogs, cats, and sheep, this region contains another four-amino acid sequence, Gly-Asn-Ser-Lys (GNSK) [4–8]. *KIT* synthesizes isoforms that differ in the presence or absence of the GNNK/GNSK sequence (GN[N/S]K).

**Fig 1.**
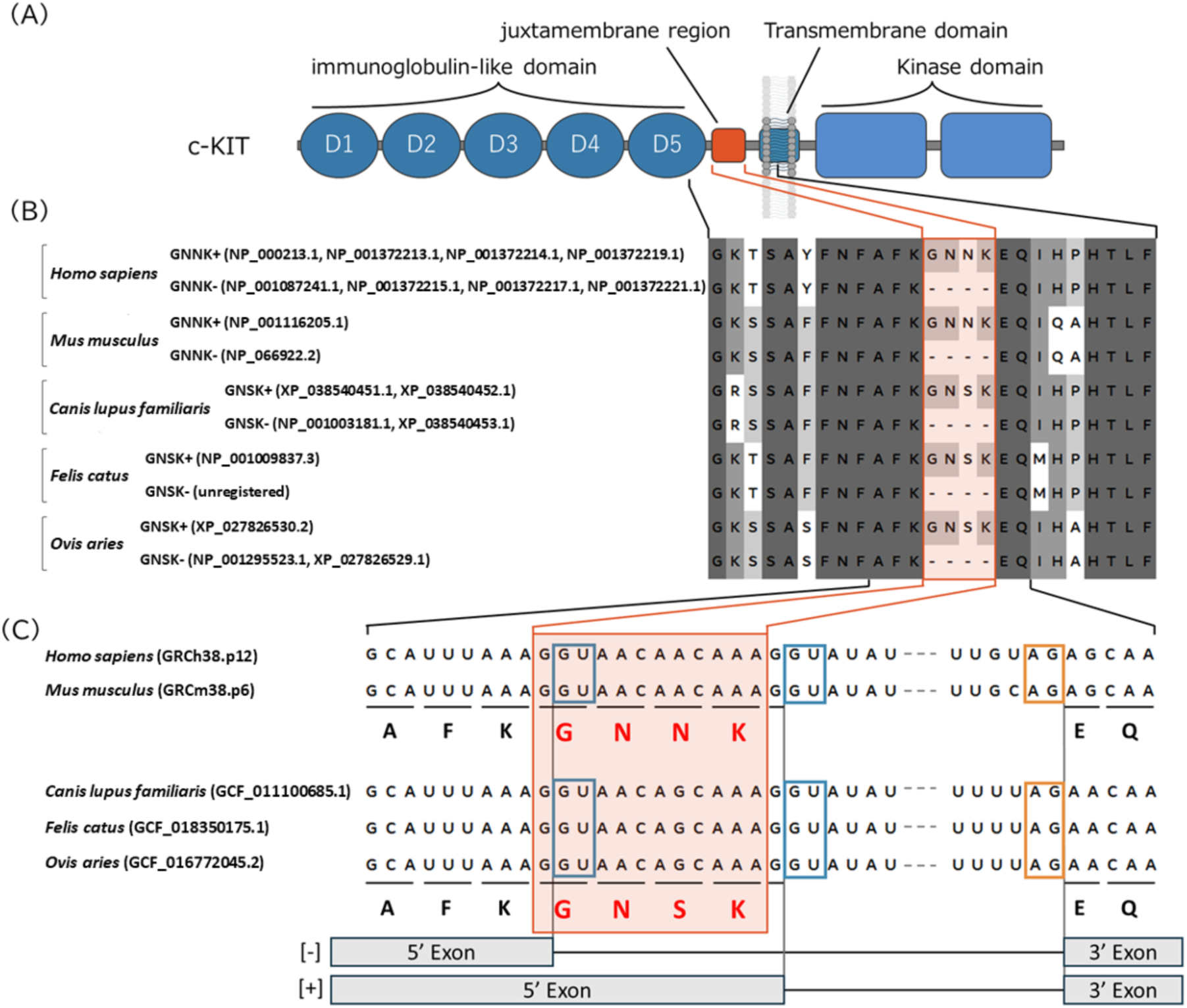
Structural features and sequences of c-KIT across five mammals. (A) Schematic representation of the structure of c-KIT protein. Blue indicates the functional domains of c-KIT, and orange indicates the location of the four amino acid sequence, GN[N/S]K. (B) Amino acid sequences around the juxtamembrane region of each species. Numbers in parentheses indicate RefSeq accession number, with prefix “NP_” representing curated Known RefSeq, and “XP_” representing computationally predicted Model RefSeq. (C) Nucleotide sequence of exon–intron boundaries around the region encoding the juxtamembrane region, arranged based on the genome assembly of each species. Blue squares indicate splice donor sites, and the yellow square indicates a splice acceptor site.

Nevertheless, there are gaps in our current understanding of these isoforms among species. In the Reference Sequence Database (RefSeq) (https://www.ncbi.nlm.nih.gov/refseq/, [9]), the isoforms with and without GNNK (GNNK+ and GNNK−, respectively) in human and mouse have been curated and registered as Known RefSeq entries, with accession numbers beginning with “NP_” (Fig 1B). In contrast, in domestic dog and sheep, only GNSK− is included in the Known RefSeq, while the GNSK+ is represented as a predicted transcript in the Model RefSeq category and indicated by accession numbers beginning with “XP_”. Nonetheless, GNSK+ and GNSK− expression in various canine tissues has been reported [5,6,10]. Moreover, the co-expression of the two isoforms in the skin tissue of Merino sheep has been observed [8]. Additionally, in domestic cats, only GNSK+ has been annotated in the Known RefSeq. However, an GNSK− isoform has also been detected in feline samples [7].

Clinically, mutations in the *KIT* proto-oncogene have been frequently reported in various cancers, including gastrointestinal stromal tumors [11,12], mast cell tumors [13,14], mastocytosis [15,16], and acute myeloid leukemia (AML) [17]. Consequently, studies on c-KIT isoforms also have mainly focused on their involvement in various diseases. For instance, in humans, the GNNK−/+ balance correlates with prognosis in AML [18]. The expression ratio of GNSK+ to GNSK− differs between healthy and tumorous canine mammary tissues [10]. Functionally, SCF-induced phosphorylation occurs via isoform-specific kinetics. GN[N/S]K− variants exhibit rapid and robust phosphorylation [7,19–23]. These differences in activity between the isoforms may be explained by differences in the orientation and flexibility of each protomer within the dimer, based on differences in the length of the juxtamembrane domain [3,23,24]. These findings suggest that isoform-dependent differences in phosphorylation and downstream signaling play critical roles in tumor initiation and progression.

Previous studies have predominantly focused on disease-specific contexts, particularly tumors or cancer, thereby limiting the range of tissues examined. Thus, studies on the comprehensive tissue-dependent characterization of the expression and regulatory mechanisms of c-KIT isoforms with GN[N/S]K presence or absence are lacking, even though *KIT* is expressed throughout the animal body. Therefore, in the present study, we performed a comprehensive expression analysis of c-KIT isoforms using tissue-wide transcriptome data from humans, mice, and domestic dogs, cats, and sheep, which should provide fundamental knowledge for clinical research.

## Materials and Methods

### Ethics statement

This study utilized publicly available data from the ENCODE database and National Center for Biotechnology Information Sequence Read Archive (NCBI SRA). As this research involved the analysis of only deidentified, publicly accessible data, it did not require ethical approval. No additional informed consent was necessary as the data were already collected and made available for research purposes. We adhered to all relevant ethical guidelines for the use of public data in research.

### Processing of transcriptome data

For the analysis of human and mouse tissue-wide transcriptome data, we first downloaded the aligned RNA sequencing (RNA-seq) data files (BAM format, aligned to GRCh38.p12 and GRCm38.p6, respectively) with their corresponding transcriptome-level gene expression quantification files (TSV format) from the ENCODE Project (https://www.encodeproject.org [25,26], S1 Table). Reference genome files and gene transfer format (GTF) files for humans and mice were obtained from GENCODE, matching the versions obtained from ENCODE: *Homo sapiens* GRCh38.p12 (GENCODE Release 29; https://www.gencodegenes.org/human/release_29.html) and *Mus musculus* GRCm38.p6 (GENCODE Release M21; https://www.gencodegenes.org/mouse/release_M21.html). To obtain splice junction data (SJ.out.tab format file) for *KIT*, we extracted reads (human: chr4:54652918-54745715, mouse: chr5:75569987-75661722) from the downloaded BAM files using the samtools view command (options -h -b) of samtools (version 1.21), and then converted them to FASTQ format using the samtools fastq command (no options for single-end sequence, and options -1, -2 to split FASTQ for pair-end sequence), and the alignment of the pairs for pair-end sequence was facilitated using repair.sh from BBMap.

RNA-seq data (FASTQ format) from samples in various developmental stages in the central nervous system (CNS) (PRJEB26969 for human and PRJEB26869 for mice) and from the knockout (KO) or knockdown (KD) experiments of alternative splicing factor [27–29] (S3 Table) were downloaded from the NCBI SRA. For the analysis of tissue-wide transcriptome data, reference genomes for domestic dogs (*Canis lupus familiaris*; GCF_011100685.1), cats (*Felis catus*; GCF_018350175.1, AnAmus 1.0 [31]) and sheep (*Ovis aries*; GCF_016772045.2) were downloaded from the NCBI Datasets portal (https://www.ncbi.nlm.nih.gov/datasets/genome/). RNA-seq data from various tissues of these mammals were obtained from NCBI SRA (S1 Table). Quality control of each raw read of the RNA-seq data was performed using fastp (version 0.22.0) [32] with default parameters. Filtered reads were aligned to the reference genome using STAR (version 2.5.2b for human and mouse and 2.7.4a for dog, cat, and sheep, using the options -outSAMtype BAM SortedByCoordinate, -quantMode TranscriptomeSAM, - outSAMattributes All, -outFilterMultimapNmax 1000, -outSAMunmapped Within KeepPairs, - outSAMstrandField intronMotif) [30]. Transcript quantification was performed using RSEM (version 1.3.1, options: -estimate-rspd, -strandedness reverse, -no-bam-output) [33].

Tissue categories were assigned based on the ENCODE “human and mouse body maps.” Samples that did not conform to any predefined category were excluded. Additionally, we introduced “adipose tissue” and “embryo” as separate categories.

### Phylogenetic analysis

ClustalW alignments were performed for the full-length *KIT* gene, including introns, and for the region encompassing the alternative exon, downstream intron, and exon of all five species. A neighbor-joining tree was then constructed in MEGA11 using the Maximum Composite Likelihood model. This model was used to compute the distances, which are expressed as the number of base substitutions per site. This analysis involved five nucleotide sequences, with all ambiguous positions removed for each sequence pair (pairwise deletion option) [34].

### Quantification of splicing variants

To distinguish between the long and the short exon of *KIT*, we focused on RNA-seq reads spanning splice junctions generated by the selective use of alternative donor sites (5′ splice sites). Based on the genomic coordinates of splice sites, reads were classified as either longer or shorter exons (for c-KIT, inclusion or exclusion of GN[N/S]K), and the number of reads supporting each splice variant was counted (S1 Fig). Samples were filtered to include only those with *KIT* expression levels of ≥1 transcript per million (TPM) and a combined total of at least three junction-spanning reads, supporting either the inclusion or exclusion of the relevant domains. Using this filtering method, we obtained 334 human samples across 14 tissue types, 184 mouse samples across 13 tissue types, 34 dog samples across seven tissue types, 76 cat samples across 12 tissue types, and 197 sheep samples across 11 tissue types for subsequent analysis (S1 Table). For data from these samples, the inclusion-exclusion ratio (in-ex ratio) for the spliced domains was calculated using the following formula:

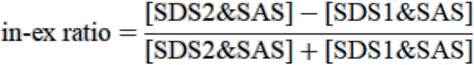

where [SDS1&SAS] indicates the number of reads aligned to the genome region containing only splice donor site 1 (SDS1) and the splice acceptor site (SAS), and [SDS2&SAS] indicates those aligned to the genome region with SDS1, SDS2, and SAS (S1 Fig). For c-KIT, [SDS2&SAS] corresponds to the count of GN[N/S]K+ and [SDS1&SAS] corresponds to that of GN[N/S]K−.

### Statistical analysis

Statistical analyses were performed using R software (version 4.4.1). Wasserstein distance was computed using the wasserstein1d function (parameter p = 1) from the “transport” package. Hierarchical clustering was computed using pheatmap function (parameter clustering_method = “complete,” scale = “none,” cluster_rows = TRUE, cluster_cols = TRUE, treeheight_row = 0) from the “pheatmap” package. Wilcoxon rank-sum test was computed using the wilcox_test function (parameter distribution = “exact”).

### Gene set enrichment analysis

The gene set files (GAF format) for humans and mice were downloaded from the Gene Ontology (GO) Consortium (https://current.geneontology.org/products/pages/downloads.html) [35,36], and those for dogs, cats, and sheep were downloaded from the NCBI portal and then converted to the GMT format. The Pearson correlation coefficient between the in-ex ratio and the expression level of all genes (TPM) was calculated using the cor function from the “StatsBase” package in Julia software (version 1.11.3). Gene set enrichment analysis (GSEA) was performed by setting the seed value with set.seed(1111) and using the GSEA function (parameter TERM2GENE={gmt files for each species}, minGSSize = 20, maxGSSize = 500, pvalueCutoff = 0.01, seed = TRUE, eps = 0) from the “clusterProfiler” package in R. Genese were ranked based on their correlation coefficient, and the top and bottom genes were used as input.

### Analysis of HITS-CLIP and IP-seq data

High-throughput sequencing-crosslinking immunoprecipitation (HITS-CLIP) datasets for NOVA2 (PRJNA286234 [37]) and RBFOX1/3 (PRJNA234443 [38]), as well as immunoprecipitation sequencing (IP-seq) data for RBFOX1 (PRJNA1134607 [39]), were downloaded from the NCBI. These FASTQ files were quality-controlled using fastp with default parameters and aligned to reference genomes using STAR (version 2.5.2b, options -outFilterMultimapNmax 1000, -outSAMattributes All, -outSAMtype BAM SortedByCoordinate). Subsequently, BAM files corresponding to biological replicates under the same condition were merged into a single BAM file using the samtools merge command with default parameters. PCR duplicate reads were removed from the merged BAM files using samtools markdup with the -r option.

### Scoring of NOVA2 binding motif

The NOVA2 binding motif (YCAY cluster) in the gene sequence was scored following Ule et al. [40]. A sliding-window analysis was performed with a 45-nt window moving by 1 nt, in which each window was examined to identify and score the following motifs.

1. YCAY followed by the next YCAY at ≥7 nt distance (YCAY-N_≧7_-YCAY) → score = 0
2. YCAY followed by the next YCAY within 3–6 nt (YCAY-N_3-6_-YCAY-N_≧7_-YCAY) → score = 1
3. YCAY followed by the next YCAY within 2 nt (YCAY-N_≤2_-YCAY-N_≧7_-YCAY) → score = 2
4. Three YCAY motifs with mixed distances (≤2 nt and 2–6 nt) being consecutive (YCAY-N_≤2_-YCAY-N_2-6_-YCAY or YCAY-N_2-6_-YCAY-N_≤2_-YCAY) → Score = 4
5. Three YCAY motifs occurring consecutively with spacing of ≤2 nt between each (YCAY-N_≤2_-YCAY-N_≤2_-YCAY) → score = 8

The total score for each window was then calculated and transformed using the common logarithm. Cross-species integrated scores were not computed; instead, species-specific scores were calculated separately.

## Results

### Two selective splicing 5′ donor sites were closely positioned in KIT

For human, mouse, dog, cat, and sheep, we confirmed that GN[N/S]K−/+ isoforms of c-KIT were generated via alternative 5′ donor site splicing that follows the canonical GU-AG rule [41,42] (Fig 1C). Within the pre-mRNA, two closely spaced GU donor sequences were present at the 5′ end of the intron immediately upstream of the exon encoding the transmembrane domain. Selection of the upstream donor site resulted in the exclusion of GN[N/S]K, whereas selection of the downstream donor site, located 12 nt downstream, led to GN[N/S]K inclusion.

### Animal species-independent tissue-specific expression of c-KIT isoforms

To clarify the whole-body expression of the isoforms, we aligned the transcriptome reads of the five species to their reference genomes to quantify *KIT* expression (S1 Table). The metadata associated with the analyzed datasets did not explicitly indicate that any samples were derived from patients with tumors or cancer. Next, referring to the splice junction coordinates, we estimated the expression of GN[N/S]K+ and GN[N/S]K−. Subsequently, the in-ex ratio was calculated for each dataset, where −1 ≤ ‘in-ex ratio’ ≤ 1 was satisfied, and ‘in-ex ratio’ = 1 when all c-KIT proteins contained GN[N/S]K, and ‘in-ex ratio’ = −1 vice versa (see Materials and Methods for details). The whole-body expression of both GN[N/S]K+ and GN[N/S]K− was identified in human (Fig 2A) and the other mammals (S2 Fig). The results revealed the following species-independent characteristics in the in-ex ratio across tissue types: Most tissues exhibited a shift toward negative in-ex ratio values, predominantly expressing GN[N/S]K− isoform. Conversely, the CNS showed a shift toward positive values, with a higher proportion of GN[N/S]K+ isoform (Fig 2B and S3 Fig).

**Fig 2.**
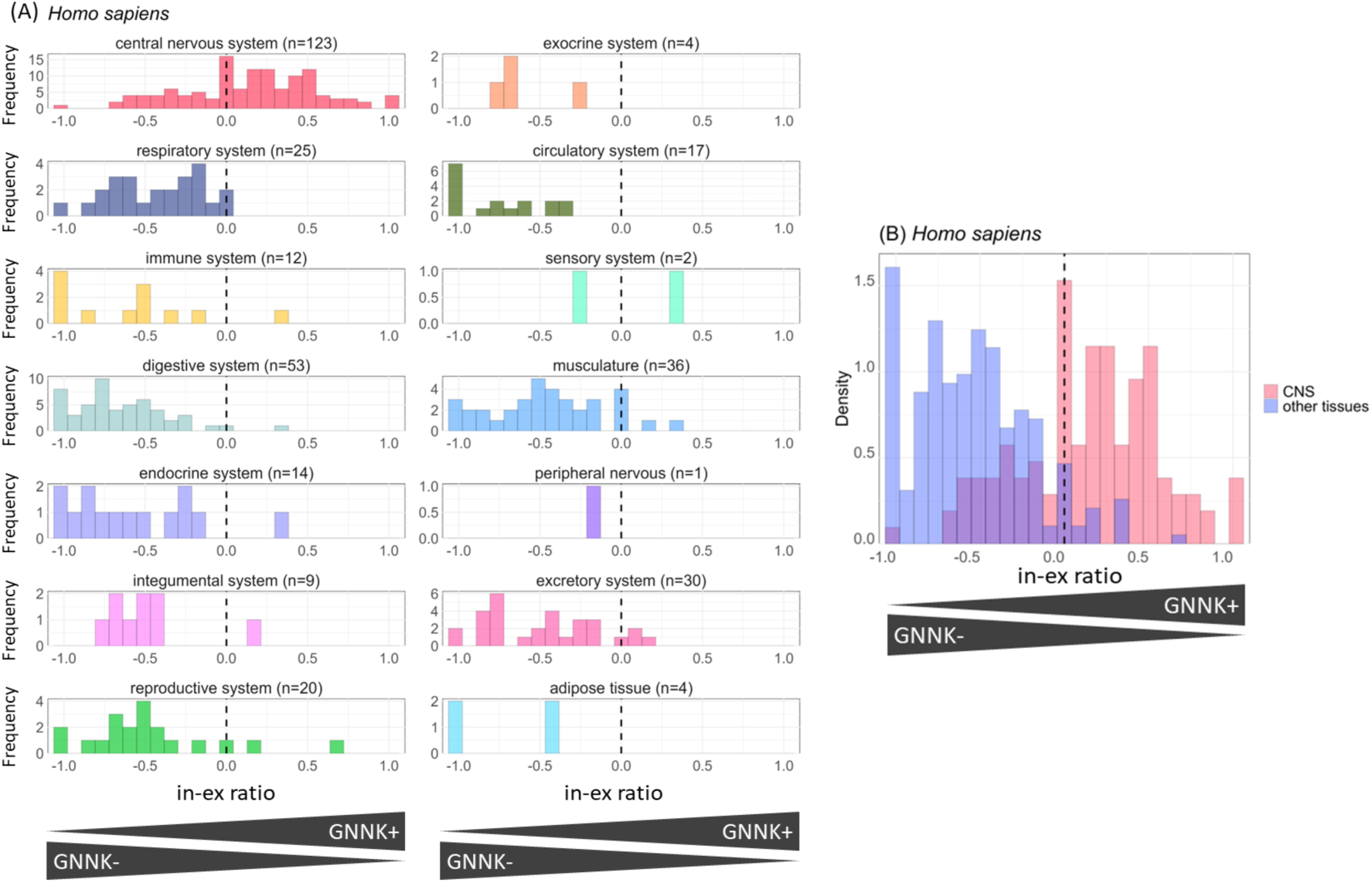
Histogram of in-ex ratio for human. (A) Histogram of in-ex ratio across 14 tissue types based on ENCODE tissue classification. “n” indicates the number of samples for each tissue. (B) Histogram of in-ex ratio colored for two categories: CNS and other tissues. Red and blue bars indicate normalized numbers of CNS samples and other tissue samples exhibiting in-ex ratio values in each bin, respectively; the total areas of red and grey bars were each assigned 1. The wavy line at x = 0 is where the expression of GNNK+ and GNNK− is equal.

To quantify the differences in in-ex ratio distributions across tissues, we calculated Wasserstein distances and performed hierarchical clustering (S4 Fig). In all species analyzed, CNS samples were consistently distinct from other tissue types in the first branch of the dendrogram, indicating that the in-ex ratio distribution in the CNS differed specifically from that in other tissues. A Wilcoxon rank-sum test revealed significant differences in the in-ex ratio values between CNS and non-CNS tissue samples: human (p < 2.22 × 10^−16^, r = 0.652), mouse (p < 2.22 × 10^−16^, r = 0.568), dog (p = 1.94 × 10^−12^, r = 0.851), cat (p = 2.95 × 10^−16^, r = 0.726), and sheep (p = 6.32 × 10^−12^, r = 0.399). For both human and mouse, the p-values reached the minimum value represented in R.

### Enrichment analysis of genes correlated with in-ex ratio

The previous section revealed the tissue-specific expression ratios of GN[N/S]K+ and GN[N/S]K− at the metadata level. To validate the tissue specificity at the gene expression level, we identified GO terms whose expression correlated with the in-ex ratio, using RNA-seq gene expression data. GSEA was performed on a ranked list of genes based on the correlation coefficient between the in-ex ratio and gene expression (TPM). In all five species, the top-ranked gene groups whose expression positively correlated with GN[N/S]K+ expression were enriched in categories related to neural development, neural function, ion transport, and cell adhesion. In contrast, the lower-ranked gene groups whose expression positively correlated with GN[N/S]K− expression were enriched in categories related to DNA repair and replication, immune response, and cell matrix (Fig 3). Consistent with these findings, and as expected, the expression of the GN[N/S]K+ isoform was strongly associated with the CNS at the gene expression level.

**Fig 3.**
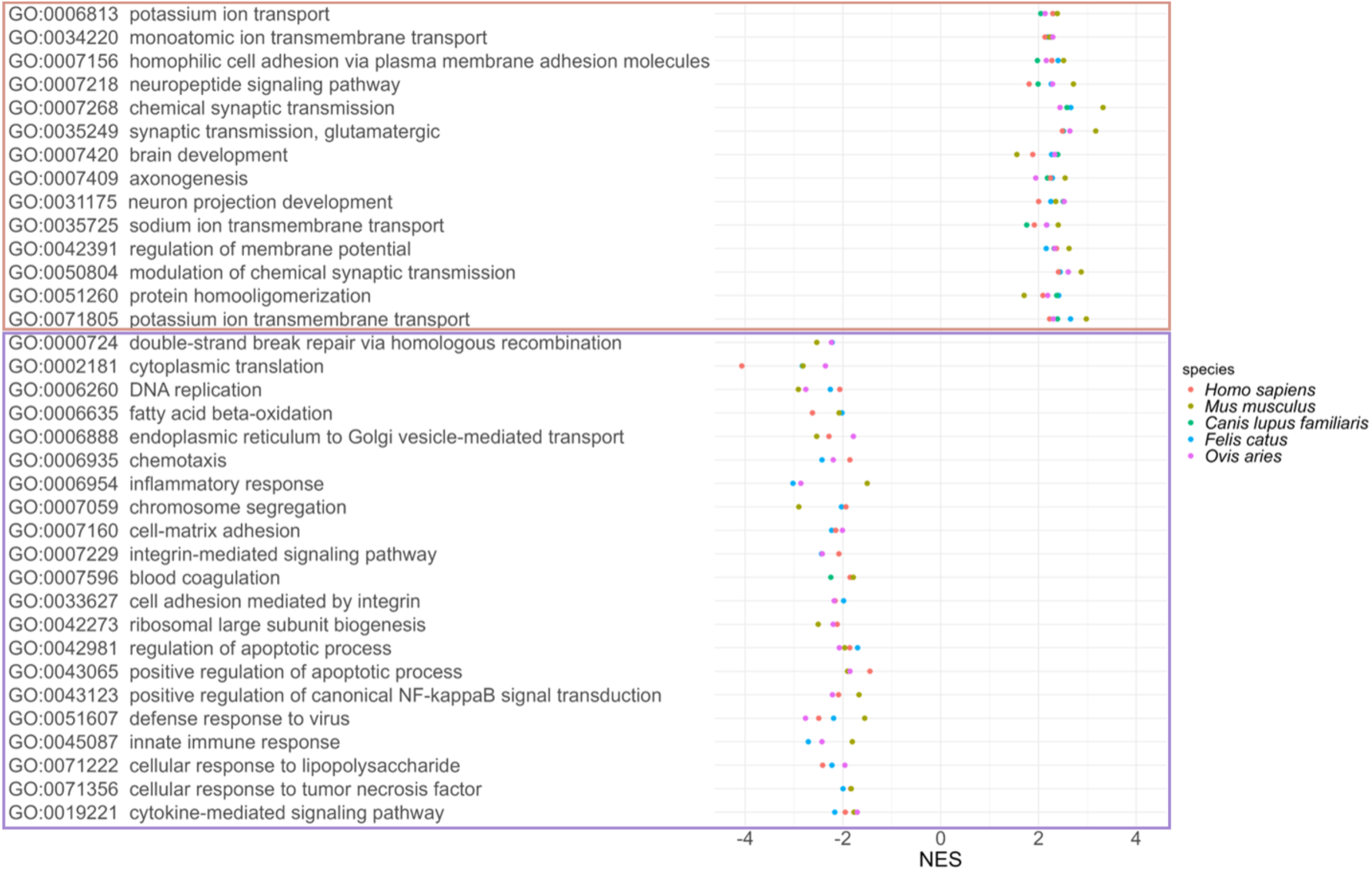
Enriched GO terms related to the ratio of c-KIT isoforms. Enrichment analysis was performed based on a list of genes sorted by the correlation coefficient between the in-ex ratio and gene expression level (TPM). Distribution of NES for GO terms and their descriptions that are shared across all five species (top). Distribution of NES for GO terms and descriptions that are common to at least three species (bottom). GO, gene ontology; TPM, transcripts per million; NES, normalized enrichment score.

### Developmental stage-dependent expression of c-KIT isoforms in CNS

Alternative splicing frequently occurs in the CNS [43] and exhibits dynamic regulation during developmental stages [44]. To capture the expression dynamics of c-KIT isoforms during CNS development, we calculated the in-ex ratio from RNA-seq data of a CNS in various developmental stages, ranging from 4 weeks post conception to adulthood in humans and from embryonic day 10.5 to adulthood in mice. Consequently, a temporal fluctuation in the in-ex ratios was identified. The GNNK− isoform predominated at early stages, whereas the relative abundance of the GNNK+ isoform increased as development proceeds (Fig 4).

**Fig 4.**
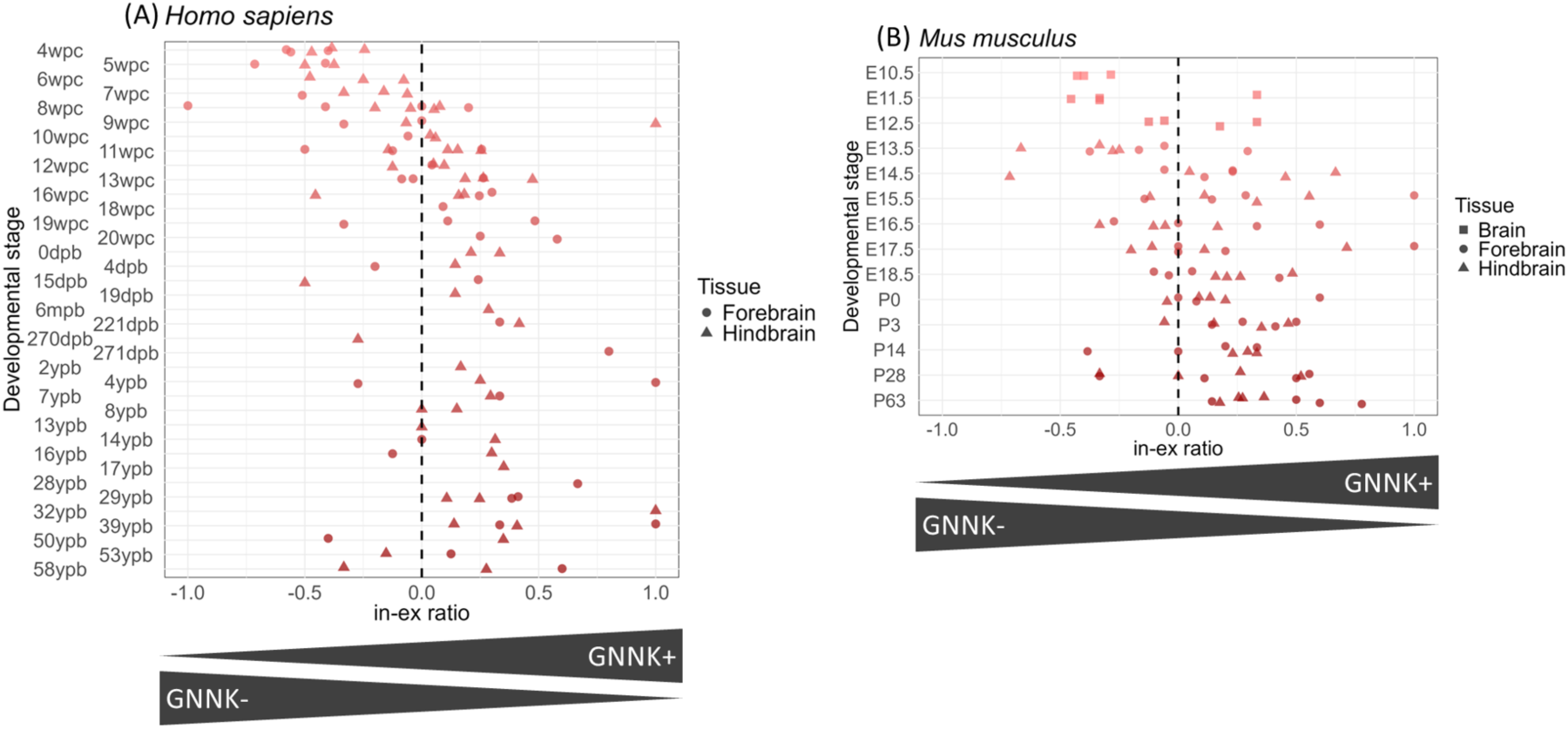
Distribution of in-ex ratio across CNS developmental time series. In-ex ratios were calculated for CNS samples from developmental time series RNA-seq datasets (human: PRJEB26969; mice: PRJEB26869). Point plots of (A) human and (B) mouse were created to depict in-ex ratios across developmental stages, with the shape of each point indicating the CNS region (human: forebrain and hindbrain; mouse: brain, forebrain, and hindbrain). wpc, week post conception; dpb, day post birth; mpb, month post birth; ypb, year post birth; E, Embryonic day; P, Postnatal day; CNS, central nervous system.

### Splice site switching is regulated by splicing factors

The expression of c-KIT isoforms was regulated in both tissue- and temporal-specific manners. We focused on genes annotated with the GO term regulation of alternative mRNA splicing via the spliceosome (GO:0000381). In a mouse tissue-wide dataset, 10 genes exhibited a correlation coefficient >0.3 (p < 0.01, FDR < 0.08) between the in–ex ratio and gene expression (S3 Table). Seven of these genes had publicly available RNA-seq datasets of KO or KD experiments, and c-KIT in–ex ratio was compared between KO/KD and wild-type samples. The datasets for *NOVA2*, *RBFOX1* & *RBFOX3*, and *DYRK1A* exhibited effect sizes that were comparable to or greater than those observed in the tissue-wide comparison analysis (Fig 5A and S3 Table). During the development of CNS, *NOVA2* expression raised sharply at embryonic day 12.5 and then declined to a constant level after postnatal development (Fig 5B). The expression of both *RBFOX1* and *RBFOX3* was activated on embryonic day 12.5 and increased monotonically. *DYRK1A*, meanwhile, exhibited little fluctuation and was consistently expressed in the CNS.

**Fig 5.**
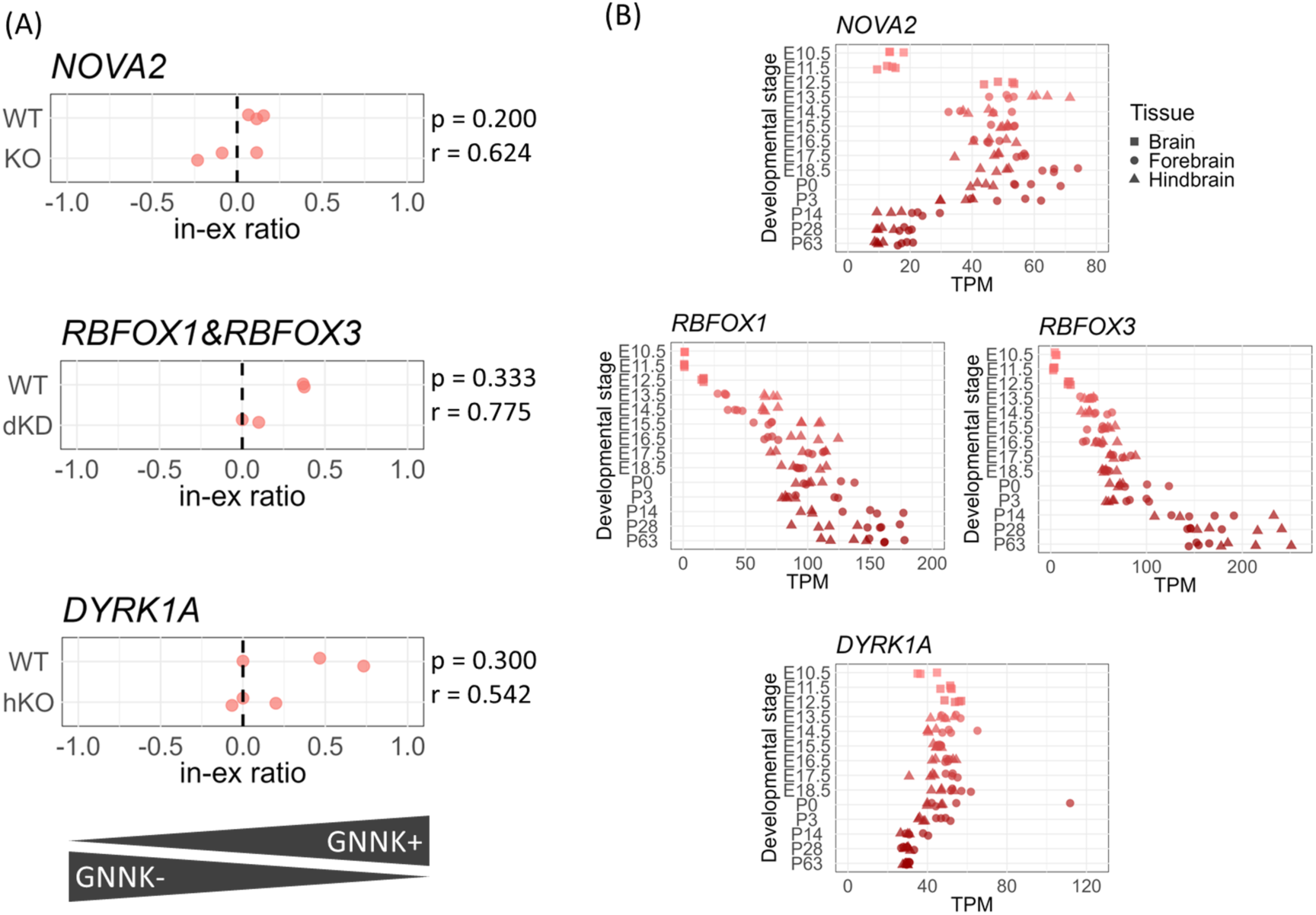
In–ex ratio distribution in *NOVA2*, *RBFOX1*, *RBFOX3*, and *DYRK1A* KO/KD experiments and gene expression across CNS developmental stages. (A) Comparison of in–ex ratio between KO/KD and WT samples: *NOVA2* KO vs. WT, *RBFOX1*/*RBFOX3* dKD vs. WT, and *DYRK1A* hKO vs.WT. “p” and “r” indicate the p-value and effect size from the Wilcoxon rank-sum test. (B) Gene expression levels of *NOVA2*, *RBFOX1*, *RBFOX3*, and *DYRK1A* across CNS developmental stages. The x-axis represents gene expression levels (TPM), and the y-axis corresponds to that in Fig 4. KO, knockout; hKO, heterozygous knockout; KD, knockdown; dKD, double knockdown; WT, wild-type; CNS, central nervous system.

### RNA-binding protein interactions and binding motifs

*NOVA2*, *RBFOX1*, and *RBFOX3* are RNA-binding proteins that regulate alternative splicing through their binding to target gene RNAs [40,45–50]. *RBFOX1* and *RBFOX3* are associated with target RNAs both directly and indirectly via the large assembly of splicing regulators (LASR) [39,51]. HITS-CLIP and IP-seq datasets were analyzed to investigate the binding of splicing regulators to *KIT* pre-mRNA.

### NOVA2

In mice, NOVA2 exhibited binding to the intronic region immediately downstream of the alternative 5′ spliced donor site (Fig 6A). We assessed the evolutionary conservation of the NOVA2-binding motif sequence (YCAY cluster) in the regions surrounding the alternative 5′ spliced donor site across species. In all species analyzed, enrichment of YCAY cluster motif was observed in the regions surrounding the alternatively spliced donor sites, where NOVA2 binding was detected in mice (Fig 6B). The RNA-binding domain of NOVA2, which consists of three KH domains, showed a high degree of amino acid sequence identity among the five species (S5 Fig). This conservation suggests that NOVA2 is likely to bind near the alternative 5′ spliced donor site in other mammals, as observed in mice.

**Fig 6.**
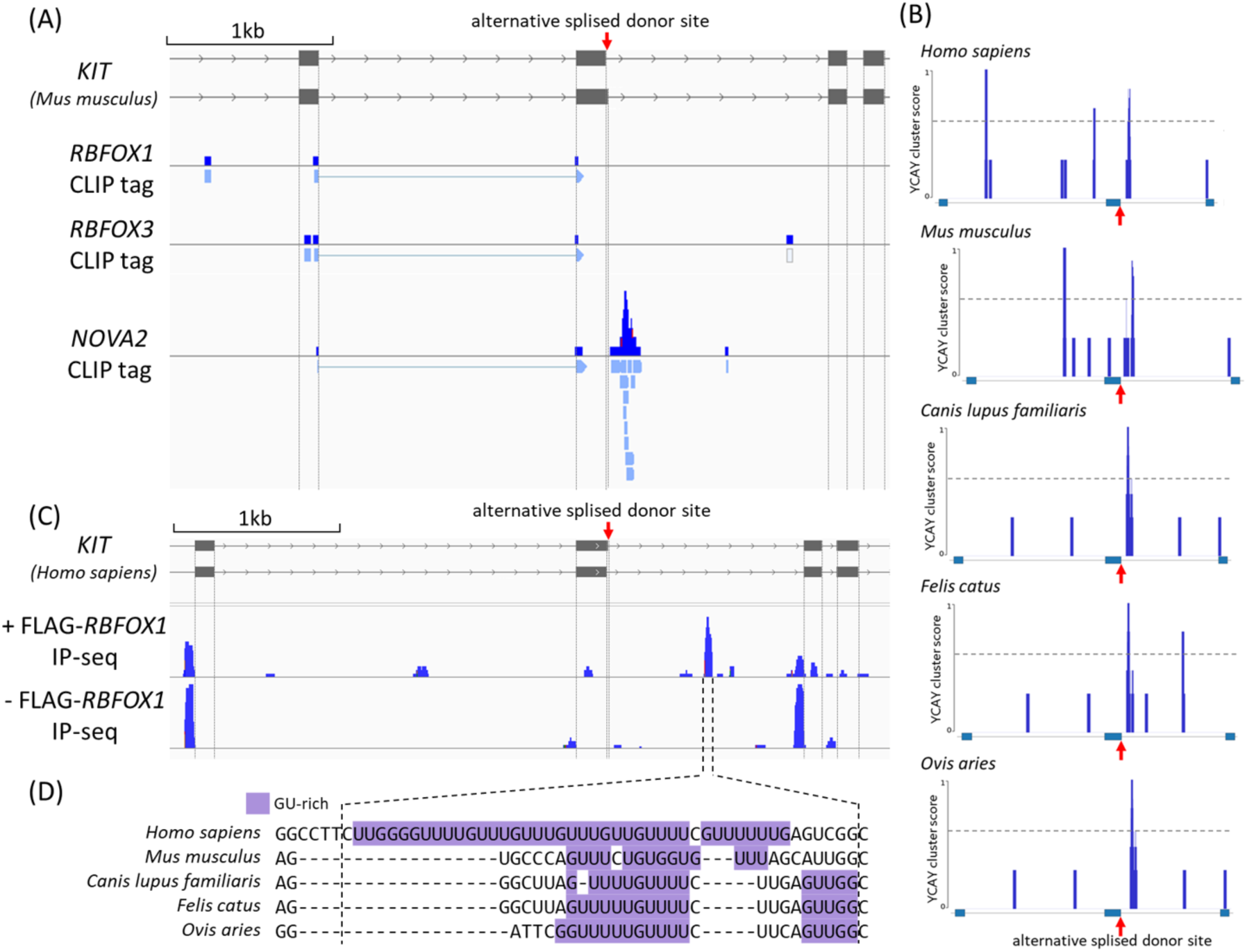
RNA binding protein interactions and motifs. (A) HITS-CLIP tags mapping for RBFOX1 (PRJNA234443), RBFOX3 (PRJNA234443), and NOVA2 (PRJNA286234) around the mouse KIT alternative 5′ spliced donor site. The figure shows sequencing read coverage (data range: 0–10) and the locations of aligned reads. The red arrow indicates the position of the alternative spliced donor site. (B) Distribution of YCAY cluster scores around the *KIT* GN[N/S]K-coding regions in reference genomes of human, mouse, dog, cat, and sheep. The horizontal dashed line represents a score of 0.6. According to Ule et al. [40], values < 0.6 represent poor conservation, or only two conserved YCAY motifs. The red arrow indicates the position of the alternative spliced donor site. (C) IP-seq tags mapping of RNAs co-purified with Rbfox1/LASR in HEK293 cells overexpressing RBFOX1. The figure shows the coverage distribution of aligned reads around the human *KIT* alternative exon (data range: 0–40). The red arrow indicates the position of the alternative spliced donor site. (D) Full-length *KIT* sequences from five species were aligned using Clustal W, and the nucleotide sequence corresponding to the region showing high coverage in (C) +FLAG-RBFOX1 IP-seq in human is shown. Purple highlights indicate GU-rich stretches (runs of four or more consecutive G or U nucleotides). Hyphens (**–**) denote deletions.

### RBFOX1 and RBFOX3

HITS-CLIP analyses of RBFOX1 and RBFOX3 in mouse revealed no enrichment of CLIP tags around the alternative 5′ spliced donor site (Fig 6A). Although a small number of single-read alignments were detected, these likely represented background noise rather than specific binding events. However, analysis of IP-seq data obtained from purified RBFOX1/LASR complexes in HEK293 cell line revealed a clear enrichment of RBFOX1 tags (+FLAG-RBFOX1) within the intron approximately 0.5 kilobases downstream of the alternative 5′ spliced donor site (Fig 6C). Conversely, no such enrichment was observed in the control sample lacking RBFOX1 expression (−FLAG-RBFOX1), suggesting that the signal was unlikely to represent background noise. IP-seq is capable of detecting not only direct RNA–protein interactions but also RNA regions that are protected by the LASR complex [39]. These results indicate that RBFOX1 associates indirectly with the intronic region downstream of the KIT alternative 5′ spliced donor site through the LASR complex rather than via direct RNA binding. In human, the sequences of the CLIP-enriched regions were prominently GU-rich, a motif recognized by the LASR subunit HNRNPM [39,52] (Fig. 6D). GU-rich elements were also observed in the corresponding regions of the other four species. This observation suggests that the LASR subunit HNRNPM may mediate direct binding to *KIT* pre-mRNA.

## Discussion

In this study, we performed a comprehensive tissue-wide analysis of the expression of the c-KIT isoforms GN[N/S]K+ and GN[N/S]K− in human, mouse, dog, cat, and sheep. Our analysis revealed that both isoforms were co-expressed in many tissues, with the GN[N/S]K− isoform generally predominant. In the CNS, the GN[N/S]K+ isoform showed a higher expression ratio, and in human and mouse, the ratio varied according to developmental stage, indicating tissue- and stage-specific expression patterns. These patterns were observed in non-tumor samples, suggesting that the expression of these isoforms exhibits tissue-specific variation independently of tumorigenesis. These observations of the dynamics of c-KIT isoforms expression are consistent with those of previous studies [4,6,7,10,53–56].

Although c-KIT is highly expressed in the brain during development and adulthood, no overt phenotypic changes were observed in *KIT* mutant mice, suggesting that c-KIT expression in the brain is functionally redundant [57]. However, subsequent studies reported that a brain-specific *KIT* mutation induced during development suppressed neural crest cell proliferation [58,59]. In addition, signaling activity differed between the GN[N/S]K+ and GN[N/S]K− isoforms through differences in phosphorylation activity following SCF stimulation [20,22]. Therefore, one potential role of the GN[N/S]K+ isoform of c-KIT in the CNS may be to fine-tune signaling activity and pathway intensity during CNS development and morphogenesis by regulating the relative expression of GN[N/S]K+ and GN[N/S]K− isoforms.

*NOVA2*, *RBFOX1*, *RBFOX3*, and *DYRK1A* were identified as candidate regulators contributing to the tissue-specific alternative splicing of *KIT*. Previous studies on *NOVA2* (and its paralog *NOVA1*) have mainly focused on the regulation of cassette exons, proposing an RNA regulatory map in which NOVA1/2 binding downstream of a target exon promotes exon inclusion, whereas binding upstream of the exon induces exon skipping [40]. Although minor alternative splicing events involving alternative 5′ donor sites, such as those observed in *KIT*, remain less well characterized, previous reports have shown that *Ccdc9* [40] and *Ncdn* [60] are regulated by NOVA1/2 and exhibit selective modulation of the alternative 5′ donor site. These findings suggest a common regulatory principle underlying NOVA1/2-mediated control, encompassing not only cassette exons but also alternative 5′ donor sites and other minor forms of alternative splicing events. Although RBFOX1 and RBFOX3 did not exhibit direct pre-mRNA binding around the alternative 5′ spliced donor site of *KIT*, RBFOX1 was indirectly associated with *KIT* intronic RNA, suggesting that the RBFOX/LASR complex may contribute to the regulation of *KIT* alternative splicing. *DYRK1A* participates in the regulation of alternative splicing by phosphorylating proline residues in the splicing factor *SRSF6* (*SRP55*), thereby modulating its activity in splice site selection [61].

The findings of this study indicate that, in both human clinical and veterinary research, it is crucial to consider tissue-specific variations in c-KIT isoform ratios when evaluating the efficacy of c-KIT-targeting agents (such as imatinib). In addition to the isoforms considered in the present study, c-KIT in sheep and cats was predicted to synthesize another type of isoform lacking the domain corresponding to exon 1 (called d1KIT in domestic cats), which might contribute to the development of the neural crest and regulation of skin coat color [8,62]. Therefore, identifying potential tissue-specific, particularly CNS-specific, functions and regulatory mechanisms of different c-KIT isoforms using in vivo and in vitro studies, as well as in silico predictions, should be a crucial focus of future research.

The present study has some limitations. The analysis of alternative splicing regulatory factors was performed primarily using data from mice (with one dataset from a human cell line included) due to the abundance of publicly available experimental resources. The correlation coefficients between the in–ex ratio and expression of regulatory factors varied among species (S3 Table), and tended to be lower in mice. This variation was likely influenced by differences in the types and numbers of tissue samples included in the datasets across species. However, in the neighbor-joining trees constructed from nucleotide alignments of the entire *KIT* locus (including introns) and from the sequence spanning the exon containing the GN[N/S]K-encoding region, the mouse *KIT* was positioned as the earliest-branching lineage among the five species examined (S6 Fig). This suggests that interspecies differences in the *KIT* gene sequence, as well as the potential presence of species-specific regulatory factors involved in *KIT* alternative splicing, may also contribute to this variation. Furthermore, owing to data availability, variations in isoform expression ratios across developmental stages of the CNS were examined only in mice and humans. Similar analyses in dogs, cats, and sheep will be necessary to clarify whether the same developmental patterns are conserved. For *RBFOX1* and *RBFOX3*, only double KD data were analyzed; therefore, it remains uncertain whether both or only one of these proteins contributes to *KIT* alternative splicing. In addition, RNA binding of *RBFOX1* mediated through the LASR has been demonstrated only in one of the human cultured cell lines, indicating that identification in vivo in humans and other species will also be required.

## Author contributions

Conceptualization: R.G. and A.A., Data curation: R.G., Formal analysis: R.G., Funding acquisition: A.A., Investigation: R.G., N.S., and A.A., Methodology: R.G. and A.A., Project administration: R.G. and A.A., Supervision: N.S. and A.A., Validation: N.S. and A.A., Visualization: R.G., Writing - original draft: R.G., Writing - review & editing: N.S. and A.A.

## Supporting information

**S1 Fig. Identification of isoforms.** Refer to the positions of the splice donor site (SDS) and splice acceptor site (SAS) in aligned RNA-seq sequence reads to distinguish between the shorter exon and the longer exon (A). Genomic locus of SDS1, SDS2, and SAS of *KIT* on the reference genome for each species (B).

**S2 Fig. Histograms of the in–ex ratio.** Histograms of in–ex ratios across tissue types based on ENCODE tissue classification. (A) Mouse (*Mus musculus*), (B) dog (*Canis lupus familiaris*), (C) cat (*Felis catus*), and (D) sheep (*Ovis aries*). “n” indicates the number of samples for each histogram. The wavy line at x = 0 is where the expression of GN[N/S]K+ and GN[N/S]K− is equal.

**S3 Fig. Histogram of in–ex ratios colored by two categories: CNS and other tissues.** The x-axis represents in–ex ratio values, and the y-axis represents the density calculated separately for CNS and other tissues. (A) Mouse, (B) dog, (C) cat, and (D) sheep. Red and blue bars indicate normalized numbers of CNS samples and other tissue samples exhibiting in–ex ratio values in each bin, respectively; the total areas of red and grey bars were each assigned 1. The wavy line at x = 0 is where the expression of GNNK+ and GNNK− is equal.

**S4 Fig. Matrix of Wasserstein distance.** Matrix of Wasserstein distance among histograms of in–ex ratios in different tissues was estimated for human (A), mouse (B), dog (C), cat (D), and sheep (E). The heatmap represents the Wasserstein distances, where each color bar range is independently scaled per species. A dendrogram was obtained by hierarchical clustering applied to a series of Wasserstein distances of each tissue.

**S5 Fig. Cross-species comparison of NOVA2 amino acid sequences.** Comparison of amino acid sequences of NOVA2 among human (NP_002507.1), mouse (NP_001025048.2), dog (XP_038278991.1), cat (XP_023100626.2), and sheep (XP_027834310.1). Sequence regions with black bars indicate KH domains. The red squares indicate positions with amino acid differences, and the numbers indicate their positions. Positions with asterisks represent identical amino acids among all five species. The amino acid sequences exhibited complete conservation among four mammals (human, dog, cat, and sheep), and mutations at seven amino acids were identified within the non-functional region of the mouse, situated between the second and third KH domains.

**S6 Fig. Phylogenetic analysis of the *KIT* in five species.** Sequence phylogenetic comparison of (A) the full-length *KIT* gene including introns and (B) the region encompassing the alternative exon, downstream intron and exon, using neighbor-joining tree method. Branch lengths above the branches represent evolutionary distances used to infer the phylogeny.

**S1 Table. Details of analyzed datasets.** (SampleInfo_targetGeneExpression.xlsx). Human and mouse: IDs of ENCODE database (experiment ID, Aligned BAM file ID, Gene expression file ID), summaries of sample, tissue type, tissue category, read count of GNNK− and GNNK+, in–ex ratio, and TPM of *KIT*. Dogs, cats, and sheep: RNA-seq accession ID, tissue type, tissue category, read count of GNSK–and GNSK+, in–ex ratio, and TPM of *KIT*.

**S2 Table. Details of GSEA results.** (GSEA_result.xlsx). Results of GSEA using a gene list ranked based on the correlation coefficient between the in–ex ratio of GN[N/S]K of the *KIT* and expression of all genes. A positive normalized enrichment score (NES) indicates that a given Gene Ontology (GO) Biological Process term is enriched among genes at the top of the ranked list, whereas a negative NES indicates enrichment among genes at the bottom. GSEA, gene set enrichment analysis.

**S3 Table. Alternative splicing factors.** (AlternativeSplicingFactor.xlsx). [corr tab] Correlation coefficients between the in–ex ratio and genes encoding alternative splicing regulators (GO:0000381) in tissue-wide RNA-seq datasets. The genes included in GO:0000381 differ among species, and if a gene is not included for a given species, its value is recorded as NA. [Mouse_experiments tab] Association between KIT isoform expression and alternative splicing regulators in mouse, based on the analysis of available experimental data for genes with correlation coefficients ≥ 0.3. The in–ex ratios of knockout or knockdown samples were compared with those of wild-type samples using the Wilcoxon rank-sum test, and p-values and effect sizes were calculated.

