## Supplementary material for "Common tissue-specific expressions and regulatory factors of c-KIT isoforms with and without GNNK and GNSK sequences across five mammals": SUPPORTING_INFORMATION.docx

**S1 Table. Details of the analyzed datasets.** Human and Mouse: IDs of ENCODE database (experiment ID, Aligned BAM file ID, Gene expression file ID), summaries of sample, tissue type, tissue category, read count of GNNK- and GNNK+, in-ex ratio, and TPM of *KIT*. Dogs, cats, and sheep: RNA-seq accession ID, tissue type, tissue category, read count of GNSK– and GNSK+, in-ex ratio, and TPM of *KIT*.

SampleInfo_targetGeneExpression.xlsx

**S2 Table. Details of the GSEA results.** Results of GSEA using a gene list ranked based on the correlation coefficient between the in-ex ratio of GN[N/S]K of the *KIT* and expression levels of all genes. A positive normalized enrichment score (NES) indicates that a given Gene Ontology (GO) Biological Process term is enriched among genes at the top of the ranked list, whereas a negative NES indicates enrichment among genes at the bottom. GSEA, gene set enrichment analysis.

GSEA_result.xlsx

**S3 Table. Alternative splicing factors.** [corr tab] Correlation coefficients between the in–ex ratio and genes encoding alternative splicing regulators (GO:0000381) in tissue-wide RNA-seq datasets. The genes included in GO:0000381 differ among species, and if a gene is not included for a given species, its value is recorded as NA. [Mouse_experiments tab] Association between KIT isoform expression and alternative splicing regulators in mouse, based on the analysis of available experimental data for genes with correlation coefficients ≥ 0.3. The in–ex ratios of knockout or knockdown samples were compared with those of wild-type samples using the Wilcoxon rank-sum test, and p-values and effect sizes were calculated.

AlternativeSplicingFactor.xlsx

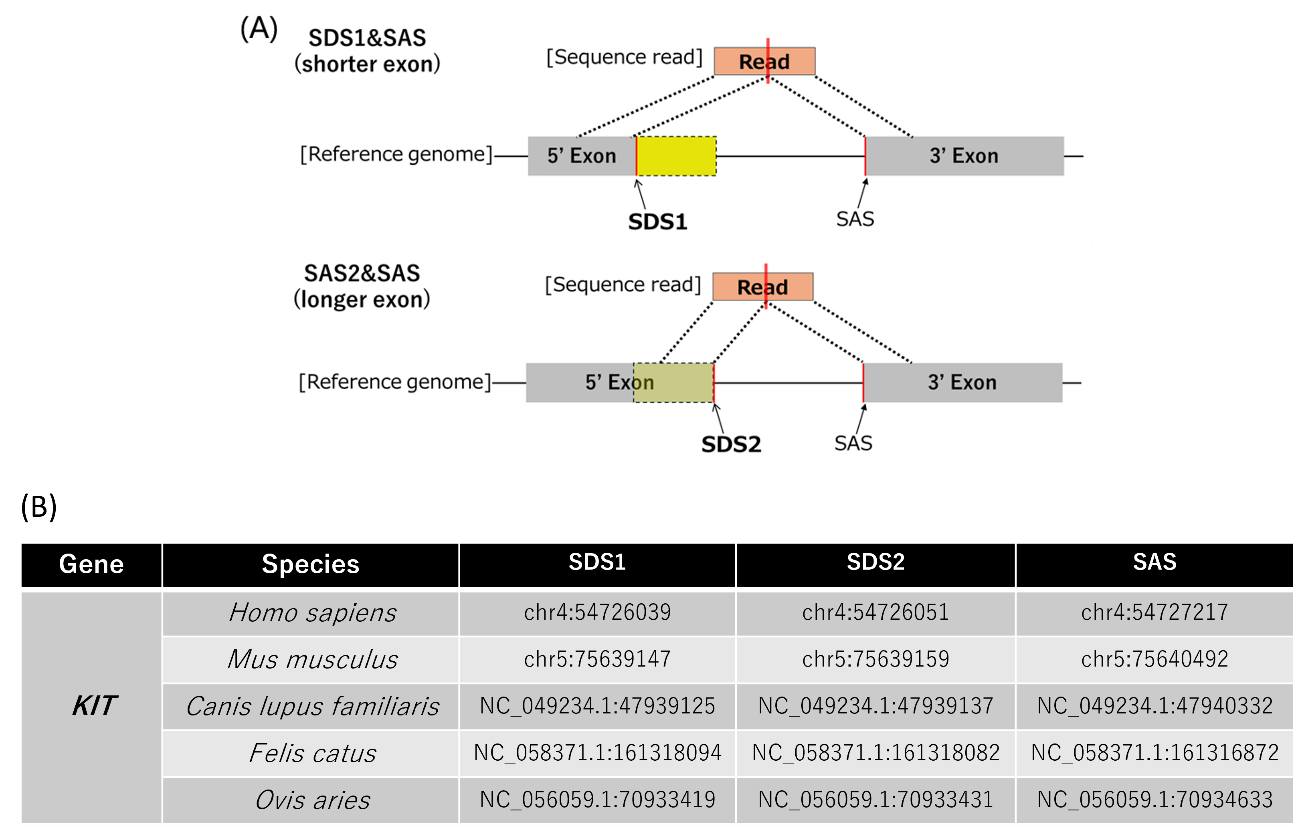

**S1 Fig. Identification of isoforms.** Refer to the positions of the splice donor site (SDS) and splice acceptor site (SAS) in aligned RNA-seq sequence reads to distinguish between the shorter exon and the longer exon (A). Genomic locus of SDS1, SDS2, and SAS of *KIT* on the reference genome for each species (B).

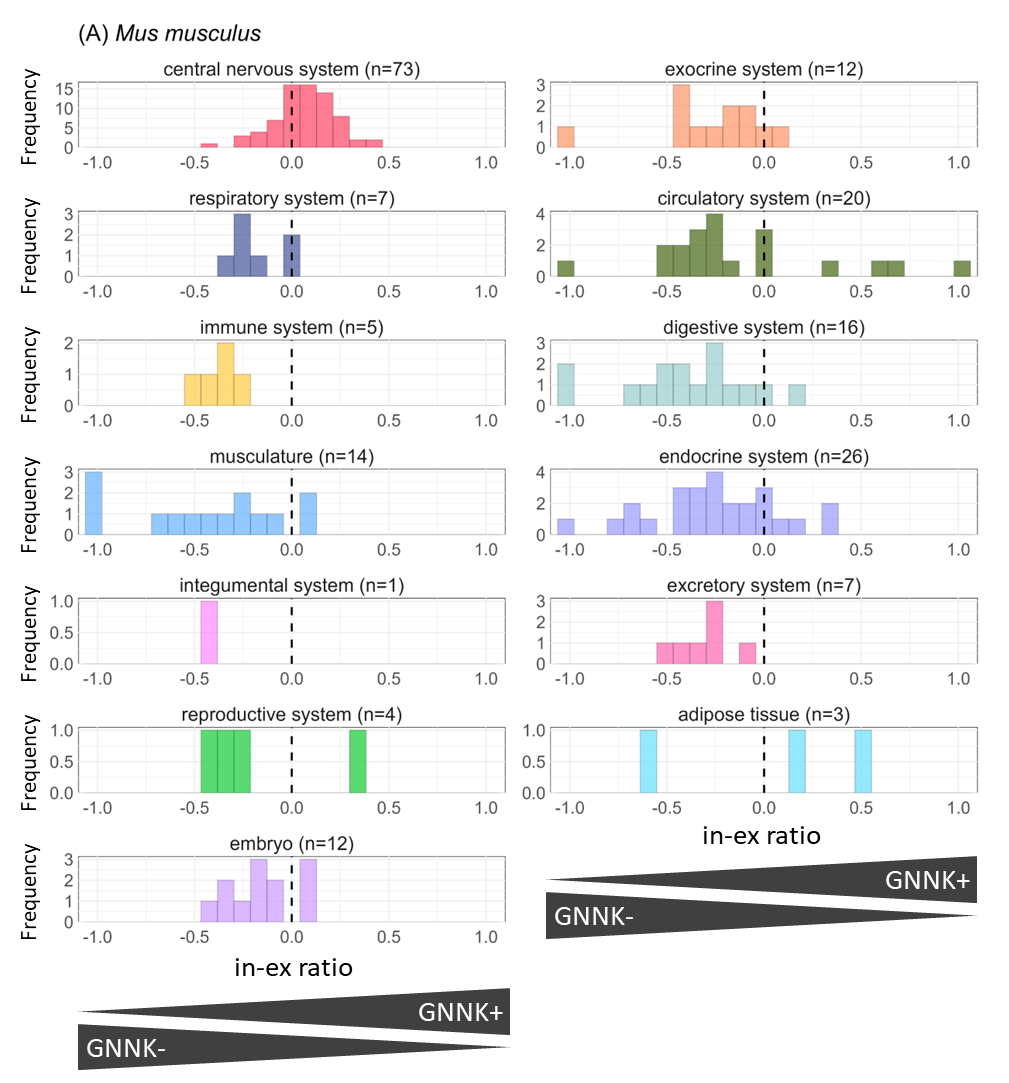

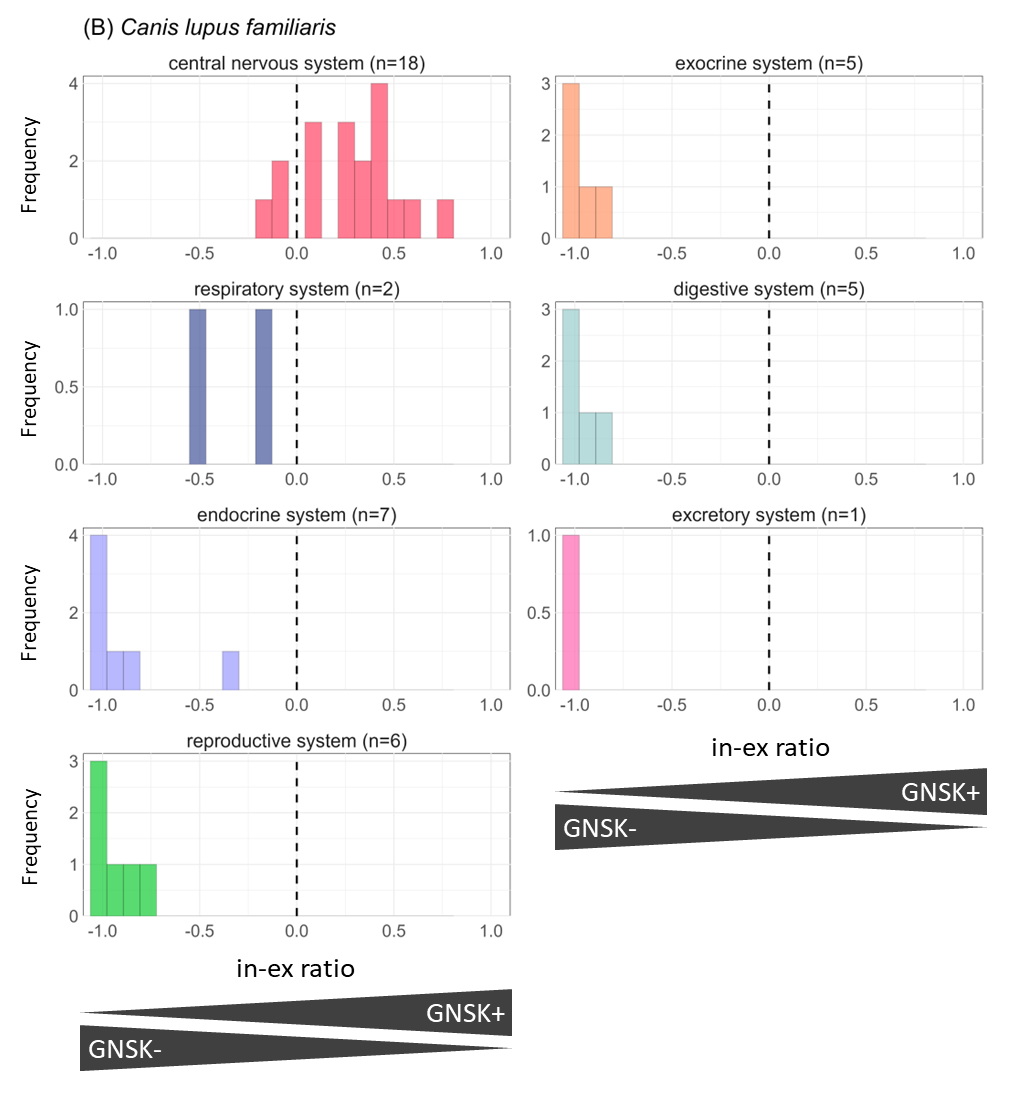

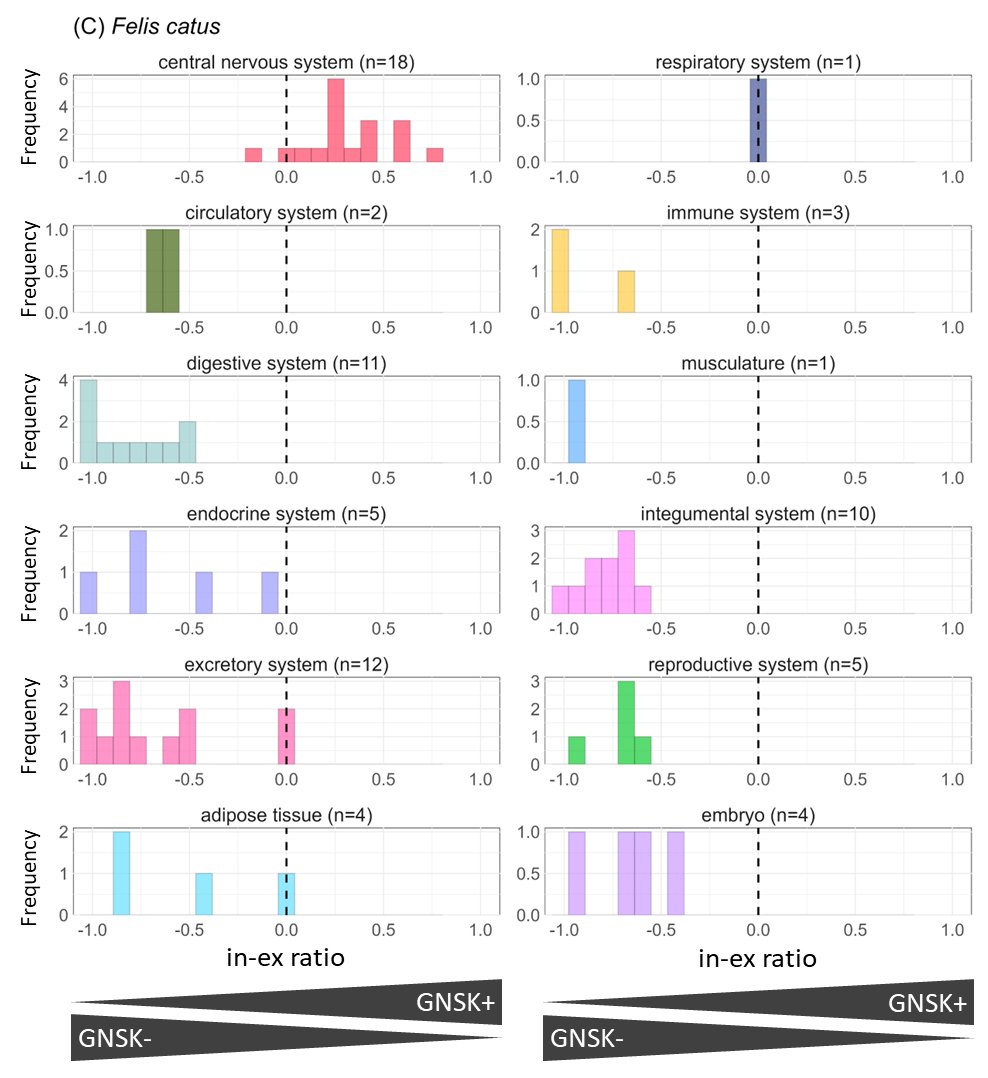

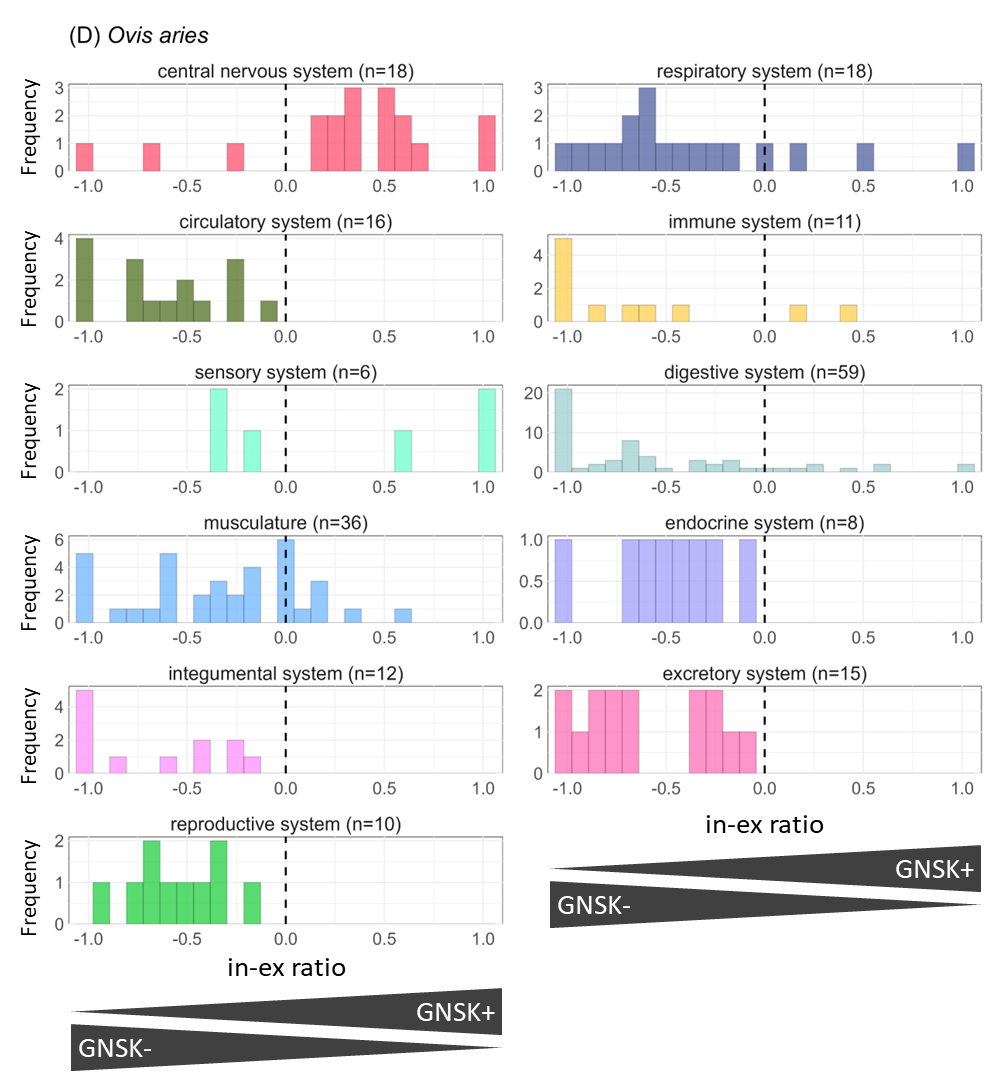

**S2 Fig. Histograms of the in-ex ratio.** Histograms of in-ex ratios across tissue types based on ENCODE tissue classification. (A) mouse (*Mus musculus*), (B) dog (*Canis lupus familiaris*), (C) cat (*Felis catus*), and (D) sheep (*Ovis aries*). The x-axis represents the in-ex ratio values, and the y-axis represents the frequency. “n” indicates the number of samples for each histogram. Draw a wavy line at x = 0, where the expression of GN[N/S]K+ and GN[N/S]K- is equal.

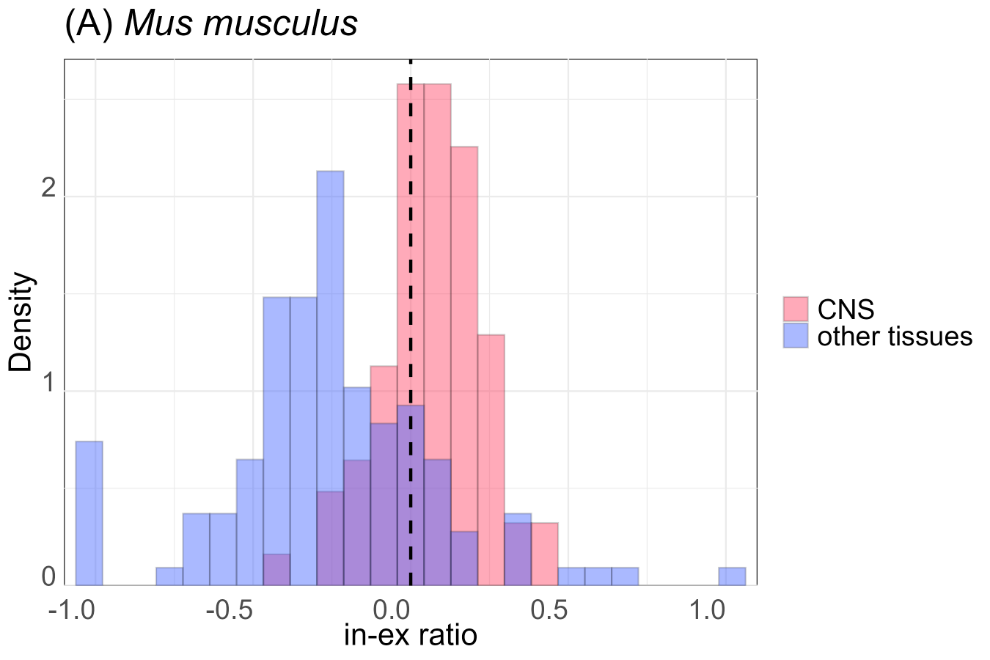

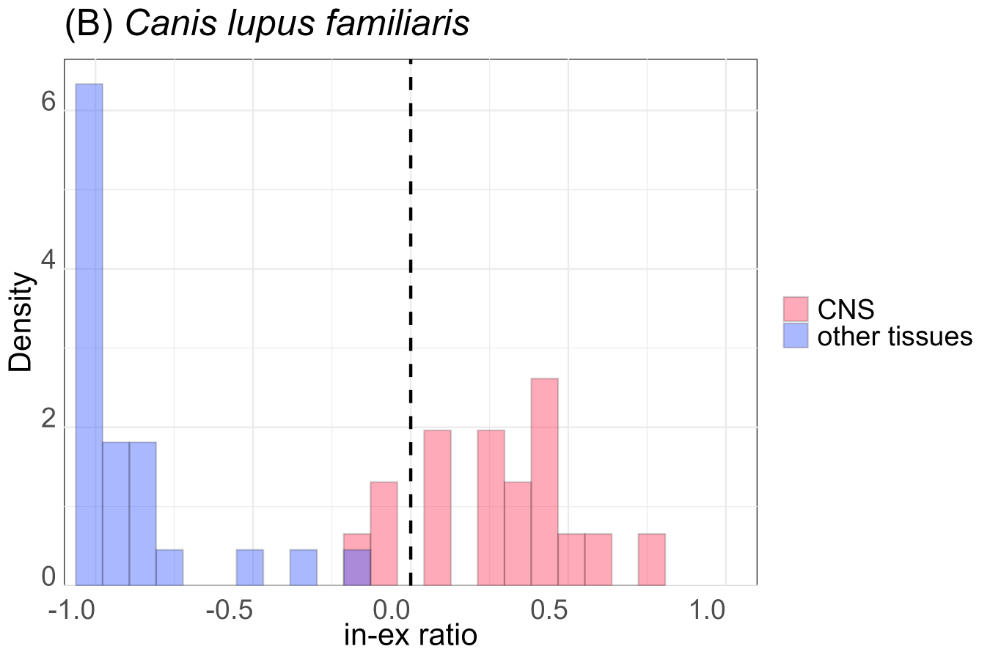

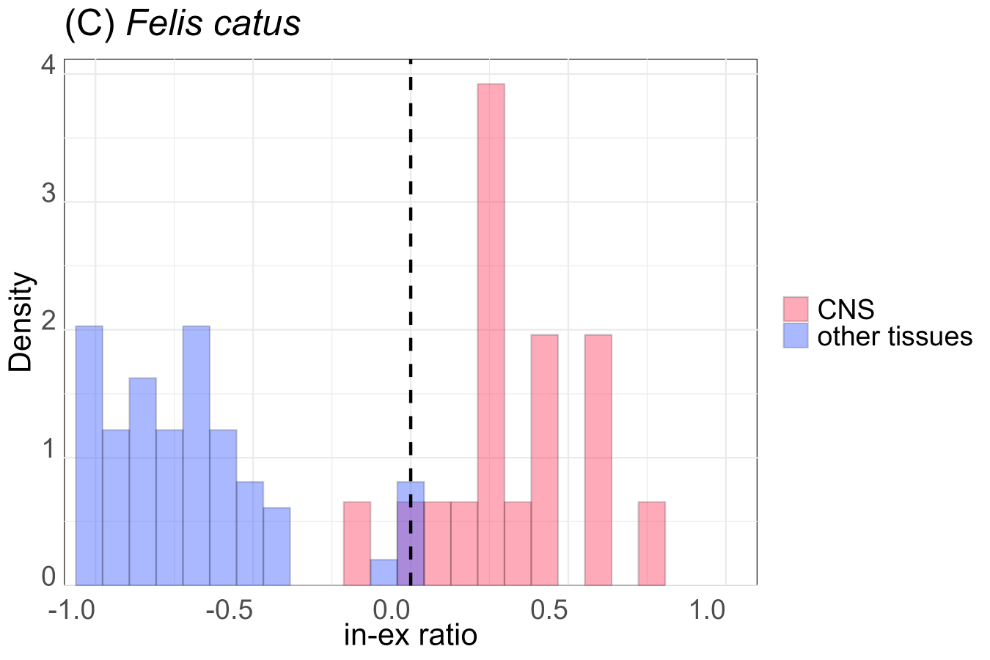

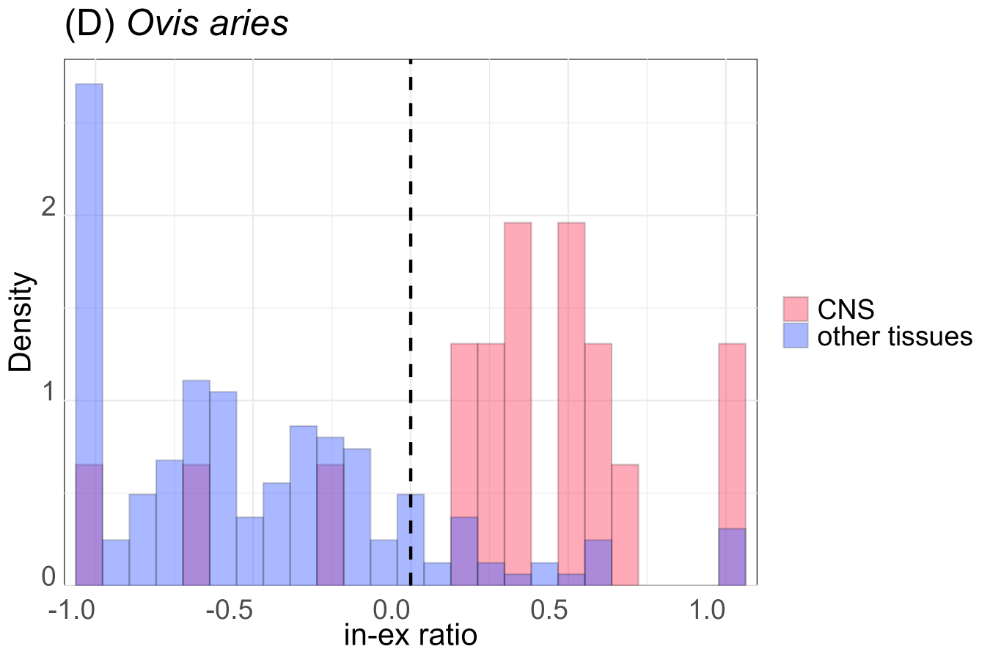

**S3 Fig. Histogram of in-ex ratios colored by two categories: CNS and other tissues.** The x-axis represents in-ex ratio values, and the y-axis represents the density calculated separately for CNS and other tissues. (A) mouse, (B) dog, (C) cat, and (D) sheep. Red and blue bars indicate normalized numbers of CNS samples and other tissue samples exhibiting in-ex ratio values in each bin, respectively; the total areas of red and grey bars were each assigned 1. Draw a wavy line at x = 0, where the expression of GNNK+ and GNNK- is equal.

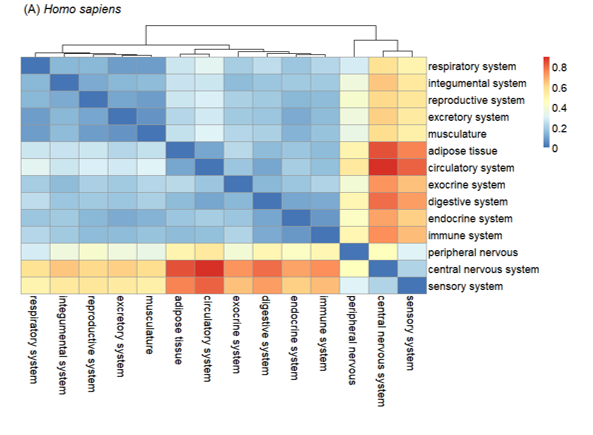

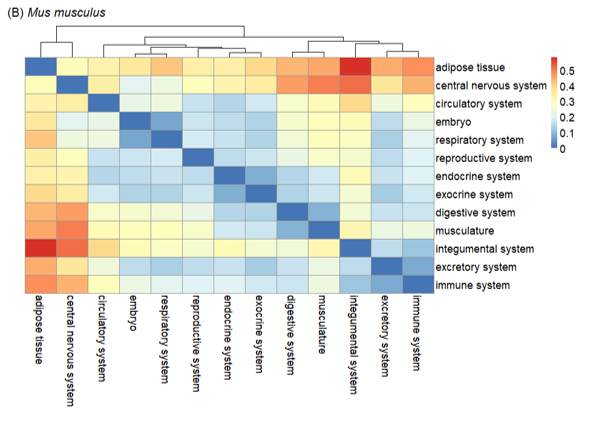

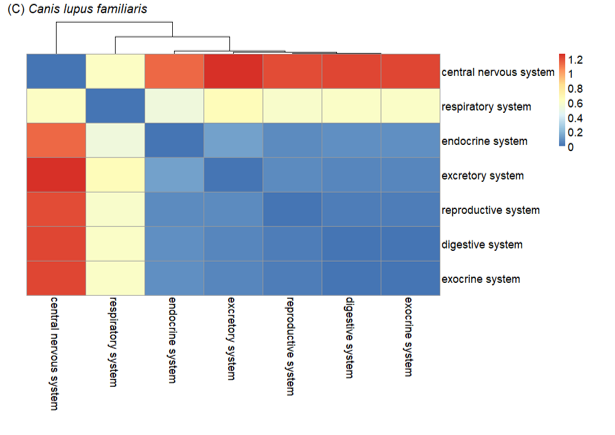

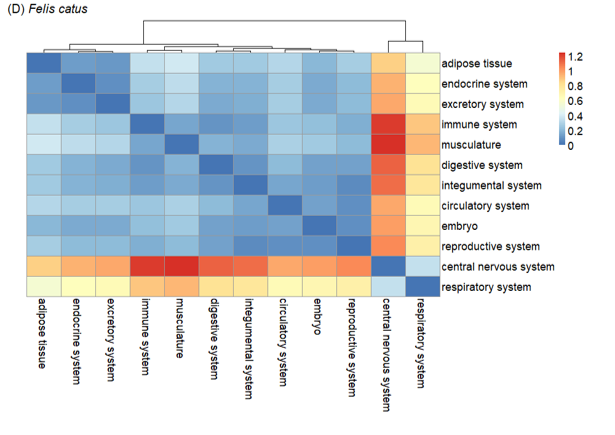

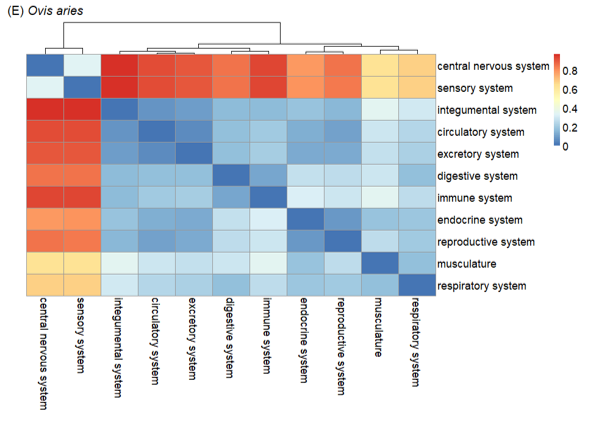

**S4 Fig. Matrix of Wasserstein distance.** Matrix of Wasserstein distance among histograms of in-ex ratios in different tissues was estimated for (A) human, (B) mouse, (C) dog, (D) cat, and (E) sheep. The heatmap represents the Wasserstein distances, where each color bar range is independently scaled per species. A dendrogram was obtained by hierarchical clustering applied to a series of Wasserstein distances of each tissue.

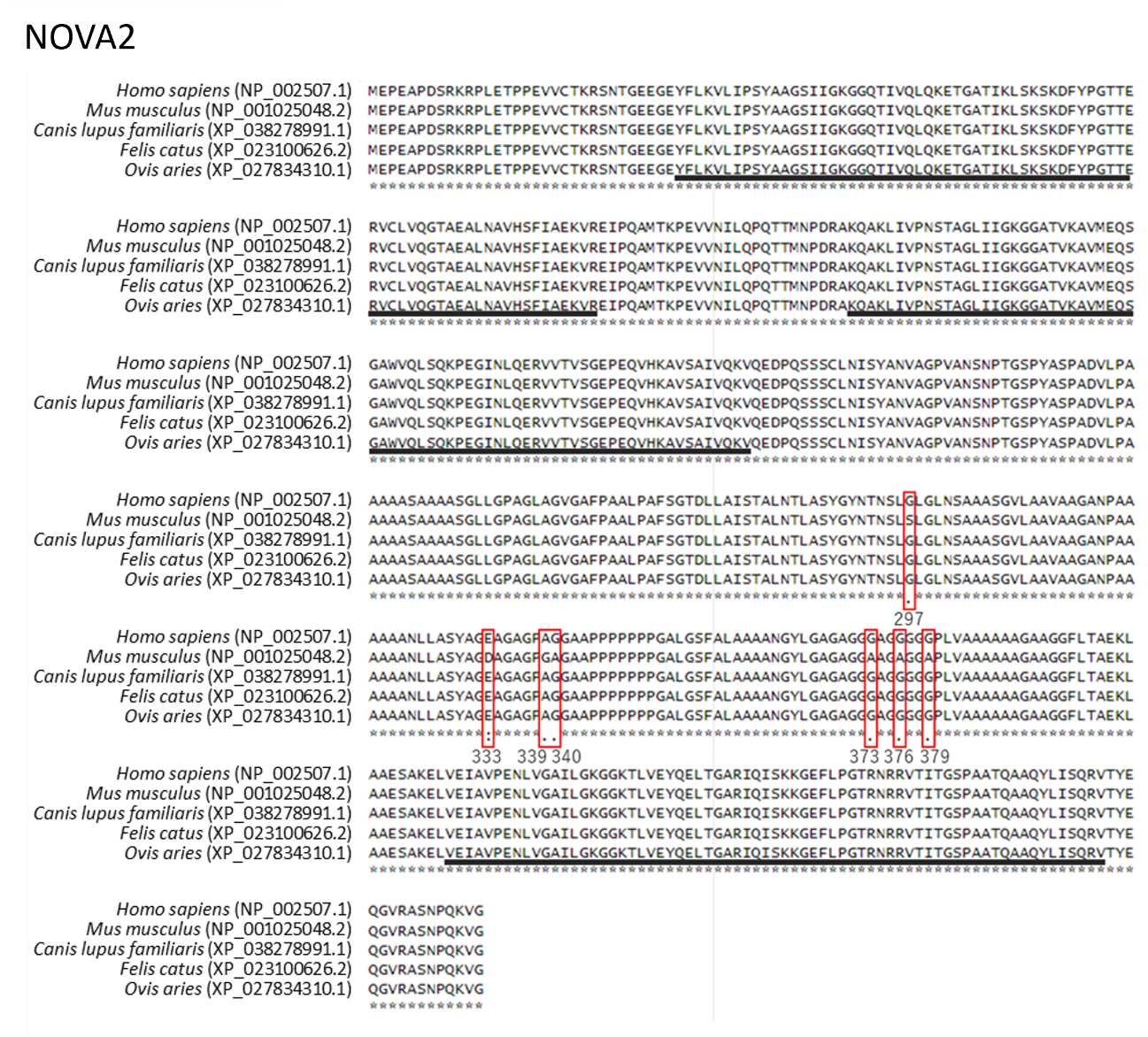

**S5 Fig. Cross-species comparison of NOVA2 amino acid sequences.** Comparison of amino acid sequences of NOVA2 among human (NP_002507.1), mouse (NP_001025048.2), dog (XP_038278991.1), cat (XP_023100626.2), and sheep (XP_027834310.1). Sequence regions with black bars indicate KH domains. The red squares indicate positions with amino acid differences, and the numbers indicate their positions. Positions with asterisks represent identical amino acids among all five species. The amino acid sequences exhibited complete conservation among four mammals (human, dog, cat, and sheep), and mutations at seven amino acids were identified within the non-functional region of the mouse, situated between the second and third KH domains.

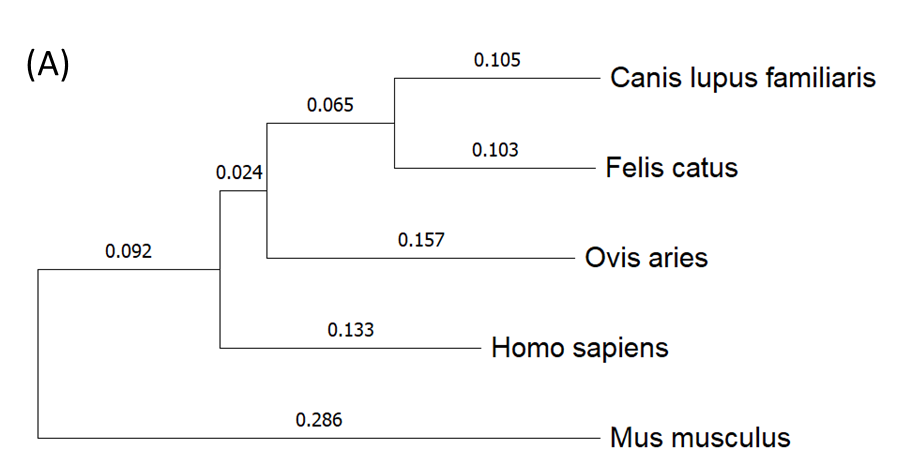

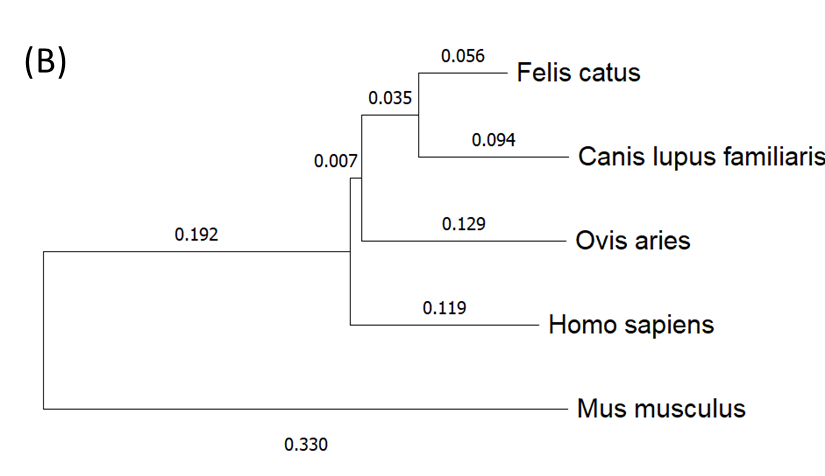

**S6 Fig. Phylogenetic analysis of the *KIT* in five species.** Sequence phylogenetic comparison of (A) the full-length *KIT* gene including introns and (B) the region encompassing the alternative exon, the downstream intron and exon, using the Neighbor-joining tree method. The values indicate genetic distances.
